# Identifying long indels in exome sequencing data of patients with intellectual disability

**DOI:** 10.1101/244756

**Authors:** Sander Pajusalu, Rolph Pfundt, Lisenka E.L.M. Vissers, Michael P. Kwint, Tiia Reimand, Katrin Õunap, Joris A. Veltman, Jayne Y. Hehir-Kwa

## Abstract

Exome sequencing is a powerful tool for detecting both single and multiple nucleotide variation genome wide. However long indels, in the size range 20 – 200bp, remain difficult to accurately detect. By assessing a set of common exonic long indels, we estimate the sensitivity of long indel detection in exome sequencing data to be 92%. To clarify the role of pathogenic long indels in patients with intellectual disability (ID), we analysed exome sequencing data from 820 patients using two variant callers, Pindel and Platypus. We identified three indels explaining the patients’ clinical phenotype by disrupting the *UBE3A*, *PGAP3* and *MECP2* genes. Comparison of different tools demonstrated the importance of both correct genotyping and annotation variants. In conclusion, specialized long indel detection can improve diagnostic yield in ID patients.

## Introduction

Whole exome sequencing (WES) continues to facilitate the genetic diagnosis of patients with intellectual disability (ID) [1,2]. The sensitivity for identifying single nucleotide variants (SNVs) from next generation sequencing (NGS) data is over 95-99% [3–5], but is lower when detecting insertions and deletions (indels), particularly in WES data (90-95%) [4,5]. Indels are known to be a key factor for human genetic disorder pathogenesis due to their damaging effect on protein structure caused by either a translational frame shift or altered protein length [6]. However, due to the difficulties in accurately detecting long indels their contribution to human genetic variation and hereditary disorders is currently under reported in many studies [7]. We aimed to clarify the contribution of pathogenic long indels in patients with ID by using specialized software targeting long indels in WES data.

The majority of SNV and indel detection methods rely on piling up reads, such as Genome Analysis Toolkit’s (GATK) UnifiedGenotyper (UG), or local reassembly on regions of interest which is the basis of GATK HaplotypeCaller (HC) [8]. Pile-up based variant callers often fail to detect long indels correctly [9] as they depend heavily on the mapping of sequencing reads and the results of alignment algorithms such as Burrows-Wheeler Aligner (BWA) [10], which apply high penalties for gaps in sequence reads for example involved in large indels. Local reassembly based methods are less dependent on prior mapping of sequence reads for variant calling and as a result have higher sensitivity and specificity in indel calling [11]. Despite their limitations, local reassembly and split-read based methods remain most suited for detecting long indels from WES data. For example, both are challenged by the fragmented nature of WES requiring both breakpoints of deletions to be inside the targeted region.

We compared two tools for detecting long indels from WES data and applied them to a cohort of 820 ID patients: 1) Pindel, a split-read targeted algorithm using a pattern growth approach [12], and 2) Platypus using local realignment of reads and local assembly [11]. Both tools can detect multiple forms of genetic variation including long indels. Pindel has previously been shown to have higher recall but lower precision rates for indels up to 100bp in size in comparison to GATK UG in WGS data [9]. However unlike Pindel, UG was not able to call indels longer than 100bp [9]. In another study Pindel had higher indel calling sensitivity for WES data if compared to GATK UG and HC [13]. However both studies note the high ratio of false-positive calls for Pindel [9,13]. Platypus has been shown to outperform other tools in short indel detection from WGS based data [14].

## Methods

Exome sequencing was performed on 820 patient-parent trios with intellectual disability [15]. Briefly, all patients were enrolled in the study from routine clinical diagnostics at the Department of Human Genetics from the Radboud University Medical Center. DNA from the patients was analysed via genomic microarrays for large CNVs as well as exome sequencing for small variants. The exomes were captured using Agilent SureSelect Human All Exon v4 kit and sequenced on an Illumina HiSeq platform with 101-bp paired-end reads to a median coverage of 75×. Mapping was performed by BWA version 0.5.9-r16 [10] and initial variant calling using GATK version 3.4 UG. We then performed variant calling using Pindel version 0.2.5b6 [12] and Platypus version 0.7.9.2 [11]. The variant calling was performed in eight batches of between 102 or 103 samples. Then the variants calls from both tools for all 820 samples were merged together using GATK CombineVariants tool and annotated with Annovar (version date 2015-06-17) [16], SnpEFF (version 4.1) [17] and VEP [18]. We focused our study on indels within size range 20-200bp. The upper limit was selected to match the median length of SureSelect v4 targets to enrich for variants with both breakpoints inside the targeted segments.

To identify clinically relevant long indels, we selected rare disruptive variants, appearing in less than 5% of the patients overlapping an ID gene list established for diagnostic WES interpretation within our department, version DG 2.3x (http://www.genomediagnosticsnijmegen.nl/services/exome-sequencing-diagnostics).

## Results

In contrast to SNV, identification the detection and annotation of longer variants such as indels remains challenging. We investigated the contribution of long indels in a cohort of patients 820 with intellectual disability, through the analysis of exome sequencing data.

We first examined the sensitivity of long indel detection of WES by selecting common exonic long indels from ethnically matched whole-genome sequencing (WGS) data derived from The Netherlands (GoNL) dataset [7] and intersecting with the SureSelect v4 targets. This resulted in 25 common (allele frequency (AF) more than 5%) exonic indels with a size range of 20-200bp and both breakpoints inside the targeted exonic regions. We detected 23 (92%) of the 25 WGS long indels using Pindel and 9 using Playpus with concordance between AFs of r=0.94 (Figure 1). Of the remaining two false negative indels, one occurred in the HLA locus with low mapping quality on chromosome 6. The second deletion had a breakpoint 9bp from the end of a capture target, probably causing the failure in detection. UG detected 17 of the 25 variants with an average 10 fold decrease in AF, and failed to detect any indels larger than 50 bp. It should be noted that although most GoNL indels were called using Pindel and Platypus, we observed a trend towards slightly lower AFs for indels identified from exome data (Figure 1). This may suggest some loss in detection sensitivity. In addition, ten more common indels occurred on the border of target capture regions and had only one breakpoint inside the exome target. In this subset, 7 of the 10 indels were identified but with a concordance between AFs of r=0.41. Resulting in a mean decrease of 70% in AF compared to the WGS AFs. In conclusion, although long indels are rare in exonic regions they can be identified using specialized software tools when both breakpoints are located within the exome capture targets.

**Figure 1.**
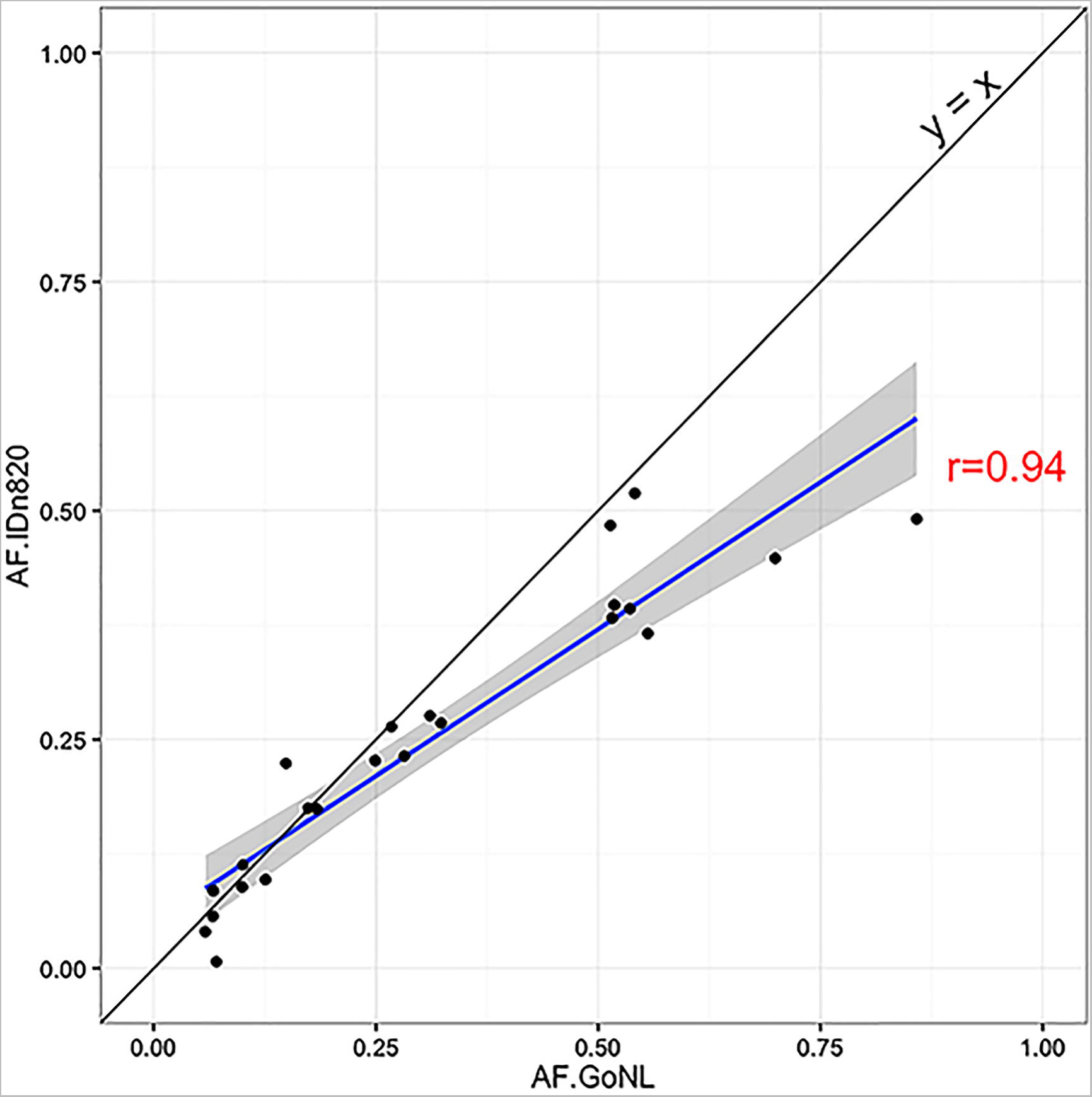
The scatter plot comparing allele frequencies for 23 common (allele frequency (AF) above 5%) indels 20-200 basepairs in size from the Genome of The Netherlands (GoNL) data having both breakpoints inside he SureSelect Human All Exon v4 (Agilent) targets compared to AFs detected from whole exome sequencing of 820 patients with intellectual disabilities. Two exons fulfilling the same criteria which appeared in GoNL data were not called in our ID patient cohort. The concordance r=0.94 is calculated using Pearson method. The linear model based smoothing indicating in blue with standard error in grey.

In total 61,052 variants were identified by Pindel and 58,443 by Platypus, of which 3,751 and 1,379 were indels occurring in the size range 20-200bp respectively (Figure 2A). Of which 92% of variants identified by Pindel and 85% for Platypus-identified appeared in less than 5% of individuals (Figure 2A). Variants were further filtered based on quality; variants identified by Pindel requiring, ten or more reads supporting variant (1,396 variants passing) and normalized Phred-scaled likelihood for homozygous-reference genotype of more than 1000 for Platypus-called variants (655 variants passing). After combining variants 1,577 unique long indels were left – 922 called by Pindel, 181 by Platypus and 474 identified by both tools (Figure 2B). 42 indels (33 in-frame, 7 frameshift, stop-gain or splice-site disrupting loss-of-function variants, and two non-exonic variants intersected with the ID gene list (Figure 2C). Many in-frame indels were located in repetitive regions, e.g. eight were detected in *FMN2* and four in *ARID1B*.

**Figure 2.**
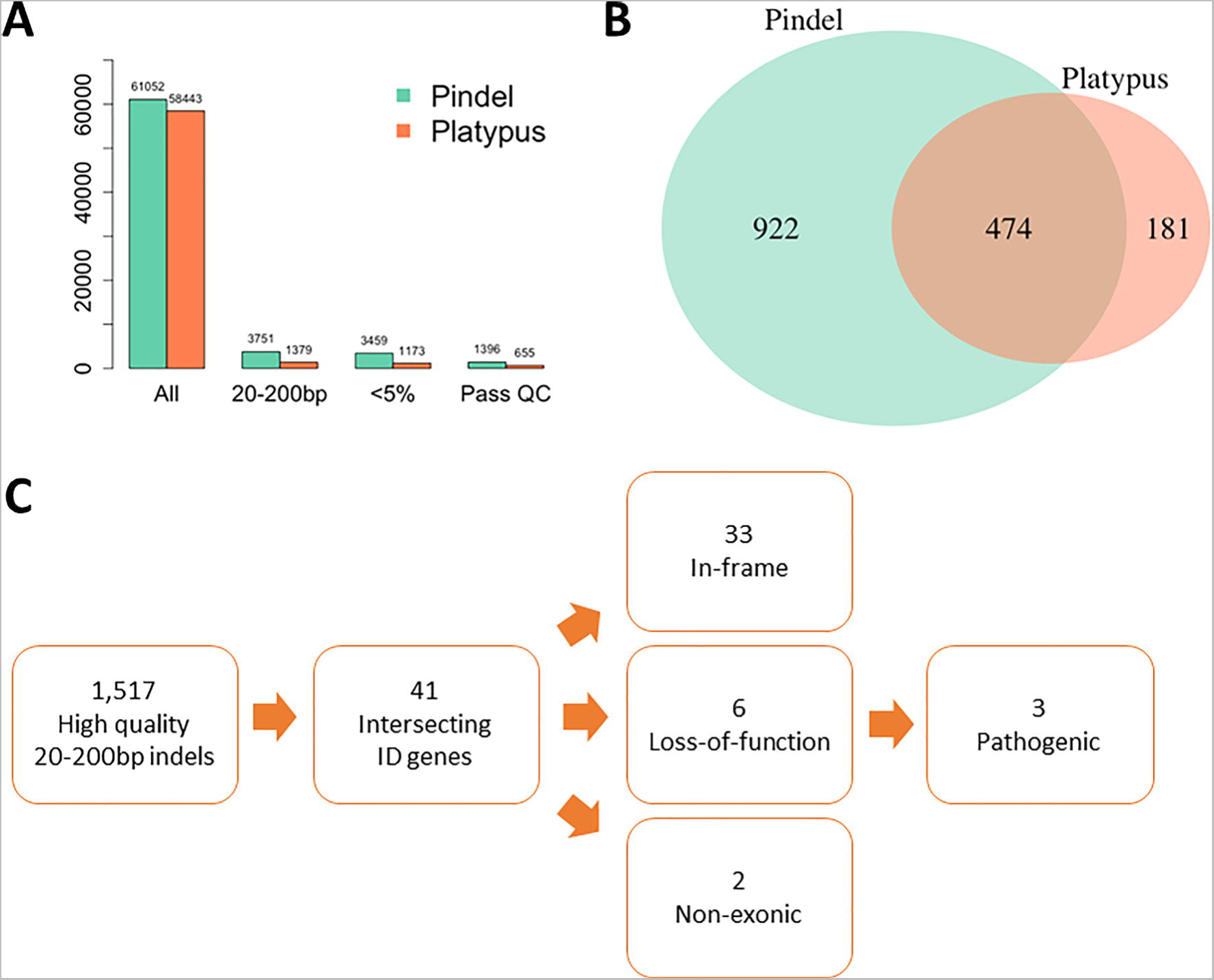
Filtering steps used for prioritizing long indels causal for of intellectual disability. A) Number of detected variants per caller after each filtering step. B) Venn diagram showing the intersection between variant callers after filtering high-quality rare 20-200bp indels C) Final filtering steps to identify probably disease-causing variants.

The seven high-quality loss-of-function variants were further assessed for pathogenicity. Two were heterozygous variants located in recessive disease causing genes *SRD5A3* and *PRODH*. However, no second damaging SNV or CNV could be identified. A 68bp insertion in the exon 10 of *CBS* gene, was detected in eight patients by Pindel, but is a known benign variant [19]. The remaining four indels represented three separate variants in *PGAP3*, *UBE3A* and *MECP2* (Table 1). All three variants were classified as pathogenic due to matches with previously reported inheritance patterns and phenotypes in the patients. All three indels were identified in the subset of 325-patients in which no causal variant had previously been identified either via SNV or CNV analysis. Thus, long indels contributed to the diagnosis of these intellectual patients in 0.37% of cases (3/820).

**Table 1.** Detected pathogenic long indels and comparison of different variant callers’ ability to detect identified pathogenic indels within the study cohort.

| Variant | Unified Genotyper (GATK) | Platypus | Pindel | Haplotype Caller (GATK) |
| --- | --- | --- | --- | --- |
| *PGAP3*:<br>c.496-39_498del<br>Homozygous | Called as heterozygous | Called as homozygous | Called as homozygous | Called as homozygous |
| *UBE3A*:<br>c.19_54delinsG<br>p.(Lys7Argfs*6)<br>Heterozygous | Not called | Called as two heterozygous variants:<br>c.19_47del and c.49_54del | Called as heterozygous | Called as two heterozygous variants:<br>c.19_47del and c.49_54del |
| *MECP2*:<br>c.1080_1194delins60<br>p.(Pro362Glufs*4)<br>Heterozygous | Not called | Not called | Called as heterozygous | Not called* |

\* Haplotype Caller called 11 heterozygous point mutations and indels up to 24bp inside the deleted *MECP2* segment.
RefSeq accession numbers: *PGAP3* NM\_033419.3, *UBE3A* NM\_130838.1, *MECP2* NM\_004992.3.

The first pathogenic indel identified was a 42bp homozygous deletion of exonintron border in *PGAP3* gene (NM_033419.3:c.496-39_498del). This deletion was detected by both Platypus and Pindel in a boy with known parental consanguinity, severe ID, epilepsy and microcephaly (Supplementary File 1). Sanger sequencing confirmed the mutation to be heterozygous in both parents and homozygous in proband as well as in an affected sib. This variant was initially detected by UG and determined to be non-pathogenic as it was genotyped as heterozygous and annotated as an inframe deletion. The initial annotation considered only the three deleted exonic nucleotides resulting in a predicted inframe deletion and not predicting the alteration in splice site.

Second, a heterozygous deletion of 36bp segment replaced by 1bp insertion predicted to cause frameshift in *UBE3A* gene (NM_130838.1:c.19_ 54delinsG p.(Lys7Argfs*6)) was called by both Pindel and Platypus (Supplementary File 2). Platypus called the variant as two separate deletions of 6 and 29bp separated by C-nucleotide matching reference, but Pindel detected the variant as a single indel. Interestingly, the maternal sample showed 5 out of 203 (2.5%) reads supporting the same deletion suggesting a mosaic deletion present in the maternal germline. This makes the maternal inheritance highly likely and consistent with Angelman syndrome matching the described patient’s phenotype. Subsequent validation with Sanger and Ion Torren PGM sequencing, confirmed the mutation both in the patient and his mother (low mosaic <10%).

Finally the third variant consisted of a 114bp heterozygous complex indel disrupting the *MECP2* gene in a girl with severe developmental delay and epilepsy, which was detected by Pindel (NM_004992.3:c.1080_1194delins60 p.(Pro362Glufs*4)) (Supplementary File 3). Both the range of deletion as well as the sequence for inserted sequence described by Pindel were confirmed by Sanger sequencing. The mutation appeared *de novo*, and thus molecular diagnosis could be established due to known X-linked dominant inheritance of *MECP2*-related disorders such as Rett syndrome.

To further assess the long indel detection capabilities of different variant callers, we performed *post hoc* analysis using GATK HaplotypeCaller (HC) (version 3.4-46) according to GATK best practice guidelines [20,21] (Table 1.) Both HC and Platypus detected the 36bp and 42bp deletions, which is expected due to the similar variant calling strategies used for both tools (i.e. local realignment and *de novo* assembly). Furthermore, both Platypus and HC called the 36bp *UBE3A* variant as two separate deletions. HC did not detect the 114bp indel in *MECP2*, but 11 *de novo* heterozygous point mutations and indels up to 24bp in size were called within the genomic segment, thus flagging the region. In contrast to the other variant detection tools, Pindel makes use of discordant read pairs in a pattern growth approach and was able to correctly detect all three pathogenic indels thus outperforming the other variant callers.

## Discussion

We analysed exome sequencing data derived from 820 patients with intellectual disability and identified 3 clinically relevant long indels which explained the patient’s phenotype. However, the application of different variant callers and annotators highlighted both the difficulties and importance of correct genotyping and annotation of long indels. Frequent discordance between annotators is well described, with splice site mutations showing the greatest discrepancy [22,23]. This is illustrated by the 42bp homozygous deletion in *PGAP3* which disrupts a splicing site. The variant was called in initial dataset, but was falsely genotyped as heterozygous variant and, moreover, annotated as an inframe exonic deletion. During this study we used two annotators – Annovar (version date 2015-06-17) [16] and SnpEFF (version 4.1) [17]. Additionally, VEP [18] was used *post hoc* to test the annotation of the *PGAP3* deletion. Annovar, also annotated the 42bp deletion as an in-frame single amino acid deletion if default arguments were used. Both SnpEFF and VEP marked the *PGAP3* indel as a high-impact variant and predicted the effect on splicing. Thus, consistent with previous publications we show the crucial role of using suitable variant annotator as well as run parameters for the variant class investigated to enable making correct conclusions on the effect and pathogenicity.

In conclusion, the disruptive indels 20-200bp in size in exons are rare cause for ID, and difficult to accurately detect using standard variant calling software. We have shown, however, that using additional variant callers and annotators can increase the diagnostic yield of WES in patients with ID.

## Acknowledgments

SP was supported by national scholarship program Kristjan Jaak (Archimedes Foundation & Ministry of Education and Research of Estonia) and Liisa Kolumbus scholarship (The University of Tartu Foundation). The Netherlands Organization for Scientific Research (NWO) has funded JHK through Veni grant 016.166.015. JV was supported by European Research Council (ERC) grant DENOVO 281964. KÕ and TR were supported by Estonian Research Council grant PUT355.

## Disclosure statement

The authors declare no conflict of interest.

